# New Genotype Emergence of DENV are Constrained by the Double Host Selection, Reducing the Probability of Alternative Serotype Emergence

**DOI:** 10.1101/639849

**Authors:** Gilberto Sánchez-González, Renaud Conde

## Abstract

Since their discovery and sequencing 40 years ago, the DENV genotypes have shown an extreme coherence regarding the serotype class they code for. Considering the RNA virus mutation rate, we used Timed Markov Model to explore the transmission possibilities of mutated viruses and the statistical eventualities of new serotype emergence. We find that around 1 000 years are required for a new serotype to emerge, in line with phylogenetic analysis of the Dengue serotypes. Our work provides a mechanistic explanation of the strictness and low probability of a new Dengue virus serotype occurrence.

**Author summary:** Recent Dengue virus global spread has drawn the attention of the Public Health Policy makers in developing countries and developed countries as well. The infection gravity and the hemorrhagic dengue syndromes have been related with the absence or presence of previous DENV immunity. Therefore, the emergence of a new DENV serotype and its spread constitute a matter of concern. Here, we constructed a mathematical model to determine the probability of such event, as well as de-entangle the mechanistic reasons behind the low serotype emergence factor of the DENV.

## Introduction

Dengue virus infection have showed a marked increase during the last two decades, imposing a sizable burden on the health system and economy of the affected Latin-American countries (1). Vaccine prospects have shown limited infection protection efficacy (42.3% to 77.7% of protection, in function of the serotype), though drastically decreasing the appearance of hemorrhagic symptoms in the infected population (2).

Part of the difficulty for Dengue vaccine design arises from the existence of 4 distinct DENV serotypes with multiples genomic sequences. Nevertheless the existence of thousands of genotypes for each serotype, the viruses have kept a great homology between the protein sequences of their structural proteins (3). This phenomenon contrast with the one observed in the human papillomavirus infection, which is caused by a DNA virus presenting over 170 serotypes (4). The occurrence of viruses not belonging to the described 4 serotypes has been reported in Malaysia, but this newcomer virus structure did not further propagate in the population (5).

Dengue virus are obliged to pass from one mammal host to one insect host to perpetuate the infection. DENV infect several human tissues, encountering cells suitable for its amplification. Typically, endothelial cells of the liver; lung vascular endothelium cells, kidney tubules multinucleated cells, spleen reactive lymphoid cells, macrophages, monocytes and lymphocytes cells are the cells suitable for Virus infection (6). In the insect, the viral particle mainly infects intestinal, circulatory and salivary gland cells (7).

In the mosquito environment, the virus mutation are dictated by the constraints of mosquito interference RNA system evasion [8], [9]. The effect of immune defense proteins, such as proteases and Thioester Protein on the virus and the mutations that would endow the virus with a better infectivity are unclear (10). In the human, the evolutionary pressure on the virus is mainly attributable to the adaptive immune system response (Ab). For instance, the E protein mutation observed between DEN-1 viruses were found primarily within the proposed immunologically reactive regions. In this particular analysis, the nucleotide variation represented only 5% of the genome, but were sufficient to evade immune pressure. All in all, this would indicate that the mutation needed for Ab reaction evasion would not require a reshaping of the serotype category epitope (11). Nevertheless, when the mutations allow a virus to escape previous immunity, the mutant usually provokes severe Dengue symptoms, including Dengue shock (12).

Dengue sole possibility to naturally persist in one host is to be transmitted vertically in the mosquito. This transmission mode has been observed in natural location (13), though has not been considered as the main route for mosquito infection. Parental mosquitoes inoculated with virus showed higher mortality than uninfected one; and when transovarially infected, the mosquitoes increased their larval stage duration and diminished their fecundity and fertility (14).

The rate of mutation rendering the virus particle inactive has been studied informatically by Burke et al. and resulted to be inversely dependent of the proposed genome length (15). One crucial aspect determining the rate of error introduced in the viruses genomes is whether they are composed of desoxyribo or ribo nucleotides. Dengue virus is a RNA virus, hence the level of errors generated during replication depends on the fidelity of its viral RNA-dependent RNA polymerase NS5. The RNA based RNA polymerase common rate of error is 1 base in 10 000 bases replicated. Indeed, a large proportion of DENV particle are defective in part of their genome, both in human patient and cultured insect cells (16). This is reflected in the differences observed between the DENV quantitation by PCR, by focus formation unit and by electron microscopy. The DENV quantities, as measured by focus forming unit, are overestimated 10 times when measured by electron microscopy and 10 000 times when assessed by PCR (17).

Interestingly, alternate passages in Huh-7 cells decrease virus fitness in Vero cells, but passage in C6/36 resulted in variable changes in fitness (increases and decreases) in Vero cells (Evolution of Dengue Virus In Vitro). Crucially, the mosquito passage of DENV impose less constraints to DENV sequence than the vertebrate, allowing them to gain diversity (18). When analyzing the field mutation rate of DENV, Khan and al. observed that the amino acid and nucleotide substitutions per year ranged from 12 to 47 and 0.35% to 1.39% of the genome, respectively. This in turn would allow for substantial variations (19). The emergence of a new serotype appears to be constrained to successive probabilistic events that would explain the resilience of the serotype classes to drastic mutation. Here, we explore mathematically the possibility of emergence and establishment of a new DENV serotype.

## Results

We first determined the number of simulations from the basal scenario whose probability of observation of an emergent new serotype was different from zero (Fig 1). From the 500 simulations only 31 resulted in new serotypes occurrence. The average probability of new serotype emergence observed in these 31 simulations was 1.1×10^-6^ [95% CI, 7.2×10^-7^ – 1.5×10^-6^] per year/per mosquito, and once averaged over the 500 simulations, 5.4×10^-7^ [95% CI, 3.2×10^-7^ – 7.5×10^-7^] per year/per mosquito. This is the expected probability of observing sufficient mutations in the sequence during a year in the mosquito population, so that a new serotype appear. This new serotype has to pass several probabilistically challenging phases until being established in the human and mosquitoes populations.

**Fig 1.**
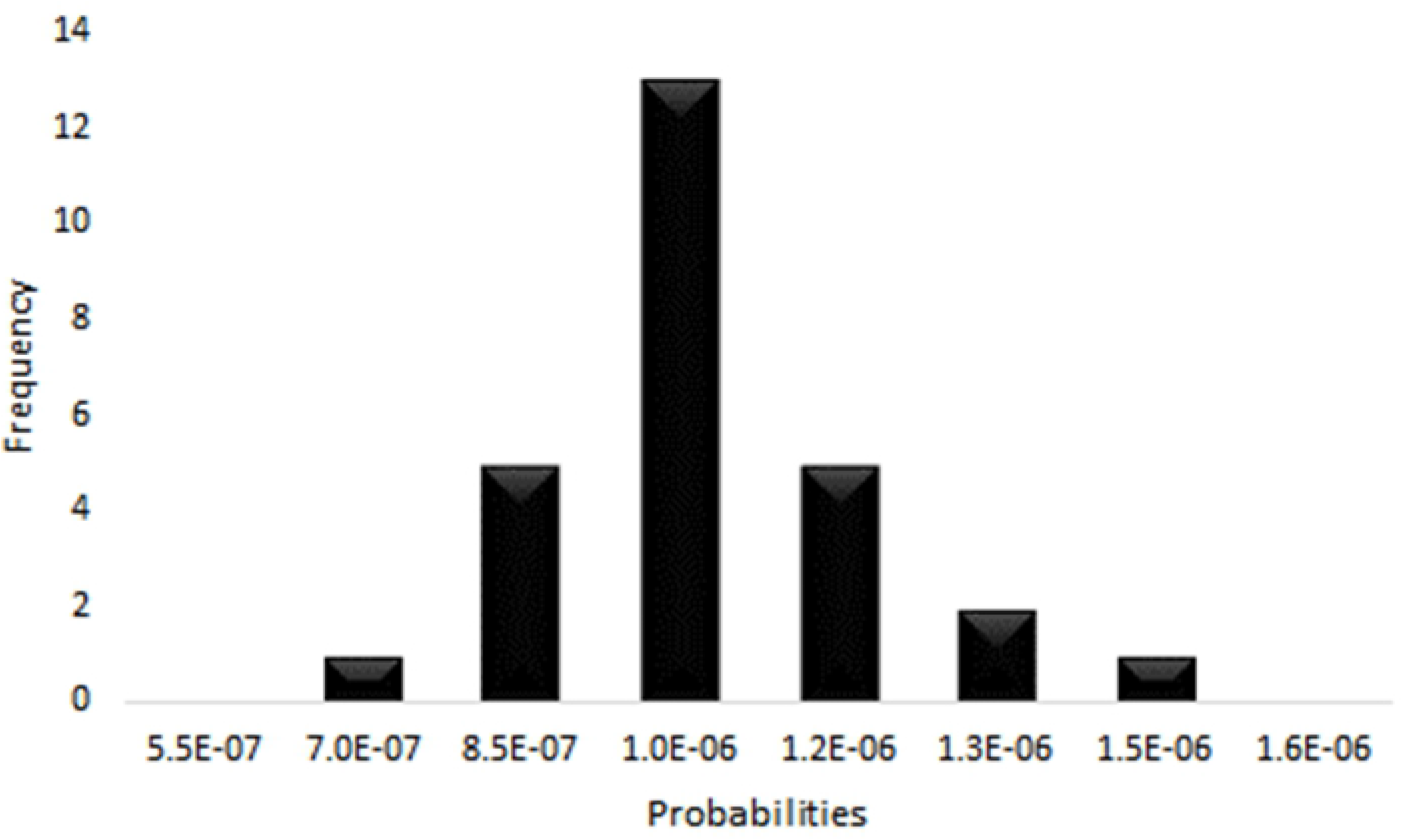
Modeling of the emergence of new Dengue serotypes for the basal scenario, were the natural mutation rate is used. From the total 500 simulations (100 000 persons and 100 000 mosquitoes each), only in 31 was observed the emergence of new serotypes.

The Fig. 2 presents the results of the simulation with vertical transmission set to zero. The column A are the results with a natural mutation rate, the column B is obtained with the natural mutation rate increased 100 times, and the column C with a natural mutation rate increased 1 000 times. The solid lines represents the mean value of these variables over the 500 simulations. The row 1 presents the scatter plots of the result of each simulation. The total human prevalence is presented with black dots. The probabilities of new serotype transmission events are presented with blue dots, and they are calculated as the number of immunological penetrations events observed in each simulation cycle, divided by the 1 000 persons simulated in each cycle. The probabilities of serotype consolidation events are presented with red dots (they are calculated in the same manner as were the new serotype transmission probabilities). The row 2 graphs the relationship between the new serotype transmission and the new serotype emergence. We observed a poor correlation between these two events. Finally, the rows 3 and 4 present the distributions of the probabilities of new serotype transmission and consolidation observed in the simulations, respectively.

**Fig 2.**
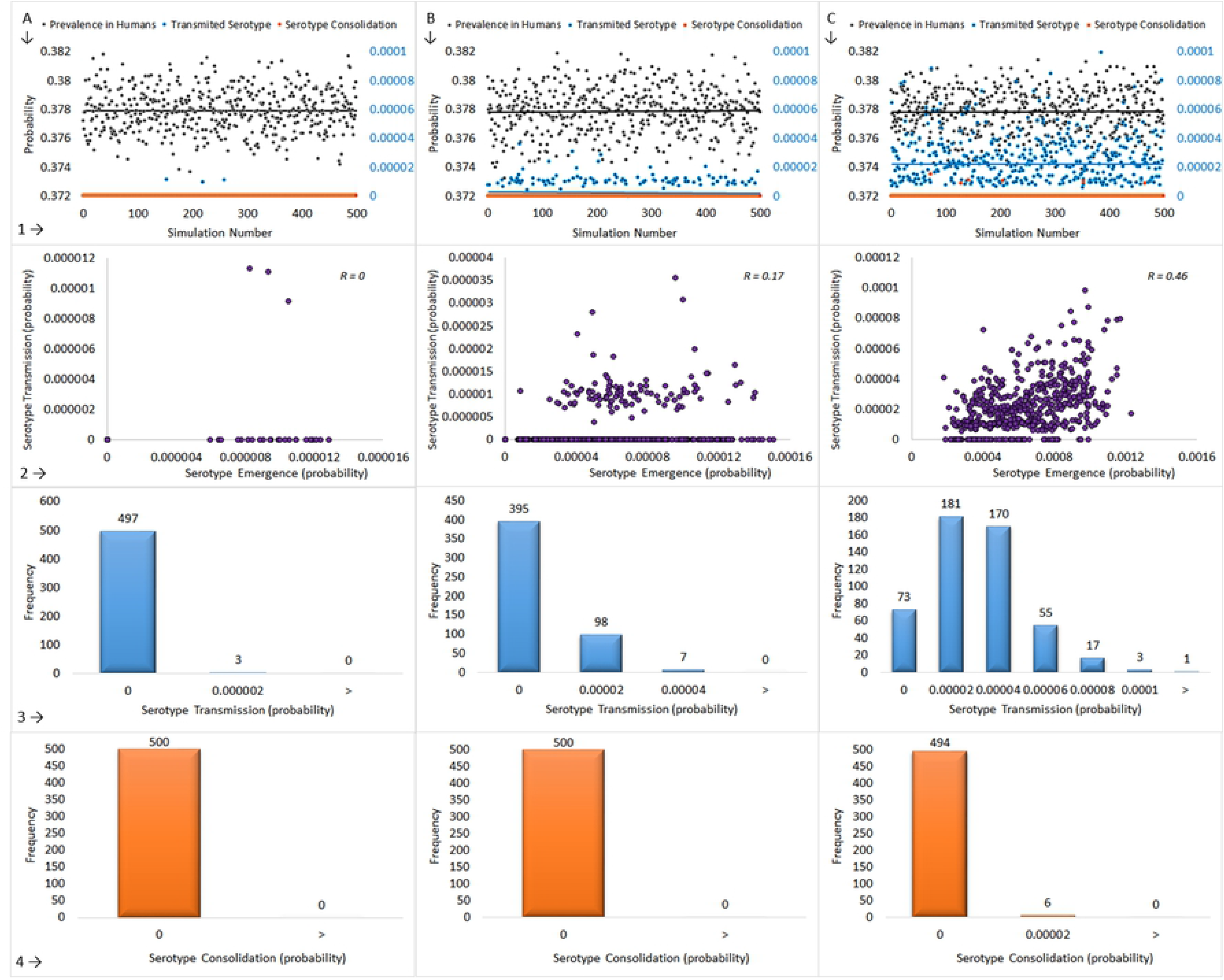
Modeling of the emergence of Dengue virus presenting a new serotype with varying mutation rate and no vertical transmission. Column A: natural mutation rate, B: 100 × natural mutation rate, C: 1 000 × natural mutation rate. The graphs represent vertically 1: 500 × (100 000 persons and 100 000 mosquitoes) simulation outcomes, 2: new serotype emergence vs. new serotype transmission, 3: serotype transmission distribution, 4: new serotype consolidation distribution.

The general description of Fig. 3 goes as follows: the emergence of sequences that satisfy our stochastic definition of “new serotype”, augmented proportionally to the increase of the mutation rate. When using 1, 100 and 1 000 times the natural mutation rate values in the equations, the new serotypes transmission to humans probabilities results were 6.0×10^-8^ [95% CI, 2.4×10^-8^ – 8.4×10^-8^] per year/per mosquito, 2.3×10^-6^ [95% CI, 1.3×10^-6^ – 3.2×10^-6^] per 100 years/per mosquito and 2.2×10^-5^ [95% CI, 1.2×10^-5^ – 3.5×10^-5^] per 1 000 years/per mosquito, respectively (when the 500 observations were averaged). This shows that, by itself, the penetration of the human immunological system represents a bottleneck for the transmission of new serotypes. This bottleneck can be explained assessing the probability of receiving a bite from a new serotype infected mosquito, multiplied by the probability of getting infected. The result derives in a minuscule final probability.

**Fig 3.**
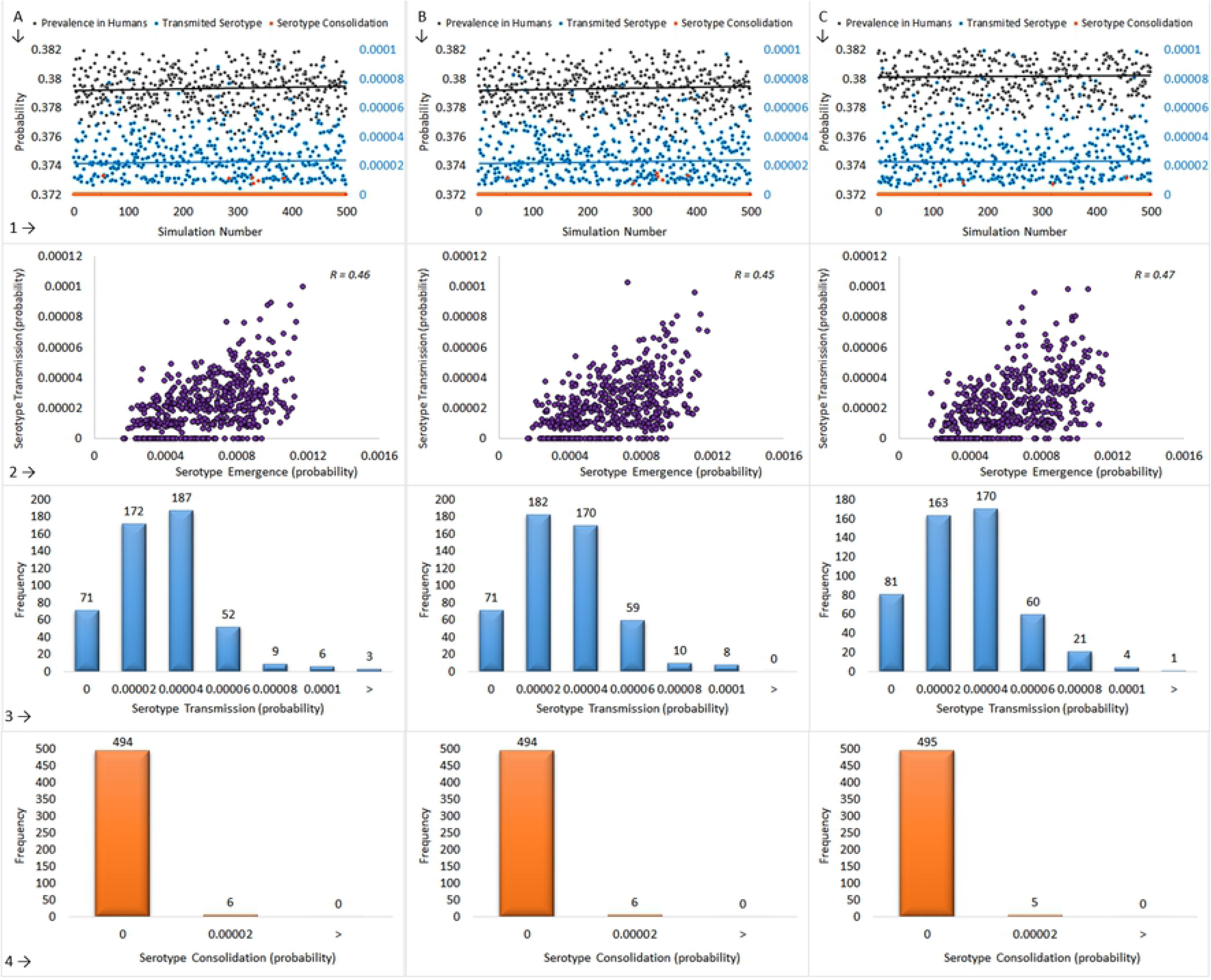
Modeling of the emergence of Dengue virus presenting a new serotype with virus mutation rate augmented 1 000 times, and with vertical transmission. Column A: 10% of vertical transmission, B: 30% of vertical transmission, and C: 50% of vertical transmission. The graphs represent vertically 1: 500 × (100 000 persons and 100 000 mosquitoes) simulation outcomes, 2: serotype emergence vs. serotype transmission, 3: serotype transmission distribution, 4: serotype consolidation distribution.

For each scenario, the serotype consolidation events resulted in a probability of occurrence of 0, 0 and 1.2×10^-7^ [95% CI, 7.2×10^-8^ – 1.7×10^-7^] per 1 000 years/per mosquito; presenting only 6 observations of probability different to zero in the 1 000 times augmented mutation rate scenario. This implies that a 1 000 years time lapse is required to obtain probabilities of consolidation of new serotypes different from zero. The hereby calculated new serotype emergence time is in line with the phylogenetic analysis of the Dengue serotypes (20).

When the vertical transmission fraction is set at 10%, 30% and 50% (Fig 3, columns A, B and C, respectively) in the 1 000 times augmented mutation rate scenario, the probability of a new serotype consolidation produces only probabilistic fluctuations, showing that this parameter is irrelevant for serotype consolidation. The transmission probability is a variable that does response to the vertical transmission increasing, but this augment seems to be insufficient for the serotype consolidation.

## Discussion

Here we modeled the mutation rate and transmission of the DENV using the transmission model in order to evaluate the serotype stability. The factors taken into account were the transition from one host to another, the RNA virus mutation rates and the probabilities of creating a new serotype/mutating the E protein in its immunogenic sequence. The model results showed that the probability of consolidation of a new serotype is particularly low, even taking into account the transovaric/transgenerational transmission of the virus in the insect vector. The model predicted that the fixation of a new serotype was only possible in a time horizon of 1 000 years, result that is in line with the reported virus serotype phylogeny time frame. This mathematical model shows that the probabilistic event of the emergence of a new serotype in the mosquito is not an odd event, with new serotype viruses appearing at a high rate. The real difficulty is its transmission to a human host and its return to the mosquito population. Hence, chance is the main driving force for the acquisition of a new serotype in the population. By this reason, we can argue that this model proposes a mechanistic explanation of the strictness and low probability of a new DENV serotype occurrence.

## Methods

A Timed Markov Model was develop to simulate the transmission dynamics between humans and mosquitoes. One of the defining attributes of Ordinary Markov Models is that the transition rates are constant and therefore the transitions are continuous. However, in our actual situation, a state transitions that depends explicitly of time is required. These are provided by the Timed Markov Models. In these models, we want to construct a continuous time process on some countable state-space *S* that satisfies the Markov property ℙ(*X_n_* = *x_n_*|*X*_*n* − 1_ = *x*_*n* − 1_), were *X* ∈ *S*, and *X* = {*X_n_*:Ω→*S*}_*n* ∈ ℕ_ denotes the stochastic process for the transition within the state-space *S*. The Timed Markov Models are defined as (21):

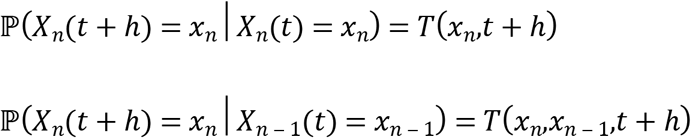

Were *T*(*x_n_,t + h*), and *T*(*x_n_,x*_*n* − 1_,*t* + *h*) are the stochastic transition rates, valuated at time *t + h.* Since in our model, the system do not express the loss of memory property (because the epidemiological state of the system can be known in any moment and thus can be compared to another similar system with different elapsed time), the stochastic transition rates *T,* cannot be (in general) an exponential distribution.

In our model, the total population of humans and mosquitoes are supposed to be in equilibria, according to the predator-prey rules (22). The model is divided in two main branches, one for humans and the other for the mosquitoes. The Humans and mosquitoes infectious rates will be evaluated from the mosquito’s branch. The human branch is split in three states that corresponds to the susceptible, the acutely infected and the immune states. In this model, the acutely infected state corresponds to the acute infection presenting high viremia levels, independently if the case is symptomatic or not.

The mosquito’s branches are divided into uninfected and infected, respectively. The infected mosquitoes first face the risk of death. If the mosquito is still alive, then the model evaluates the probability for the mosquito to produce a new serotype or to remain with the consensus sequence. Once determined this condition, the mosquito may or may not enter to a meal search period, where it will be searching for a prey to feed on. If a consensus sequence carrier mosquito bites a susceptible human, the human faces the risk of getting infected; but the bite of this type of mosquito cannot infect the acutely infected or immune persons.

If a new serotype carrier mosquito bites a susceptible human, the human faces the risk of getting infected, and this event will be considered as a “transmission of a new serotype event” (TNSE) and it will be counted by the model. These mosquitoes have the potential to modify the health state of immune persons, because this new serotype will evade their immune system and produce a new acute infection and another TNSE. Since we take into account the cross protection mechanisms effects, this new re-infection will be only possible when the infected human has no cross-serotype protection. For the same reason; in this model, an acutely infected person cannot be re-infected.

On the other hand, at an initial point, an uninfected mosquito also faces the risk of death. If the mosquito is still alive, then the model evaluates the probability, for the mosquito, to enter to the meal period. If the mosquito bites a person infected with a consensus sequence, the mosquito faces the risk of getting infected. Similarly, if the mosquito bites a person infected with a new serotype, the mosquito risks to get the infection and this event will be counted as a “new serotype consolidation event” (NSCE). The incidence of the disease in the humans determines the probability for the mosquito to bite an infected person. The size of the human incidence is determined at each time step of the model. Over time, the accumulation of new cases increases the DENV incidence. In the model, for each mosquito that dies, a newborn mosquito emerges. This new mosquito will be naïve or infected born, depending on the scenario (with or without transovaric transmission).

In the human’s branch, the susceptible persons remain susceptible until they are infected from the mosquito’s branch. If they are infected, they enter the acute period, where they remain some period of time, until they probabilistically enter the immune state. This state can be abandoned losing the long-term immunological protection (and returning to the susceptible state) or by reinfection with a new serotype (and returning to the acute infection state).

The model evaluates the states of the humans and the mosquitoes concurrently. To produce the alternated changes between the infected populations, we force the model to enter the human branch once the mosquito branch has been evaluated. This allows to achieve a human state response to the mosquito state at the same step of time. The time step size of the model is a day. The model evaluates and averages the results of simulating the infection of 100 000 humans and 100 000 mosquitoes during a rain season (4 months modeled as a sine-square function). This procedure is repeated 500 times, with different parameters values obtained from a random sampling of probabilistic distributions (see the model scheme at Fig 4).

**Fig 4.**
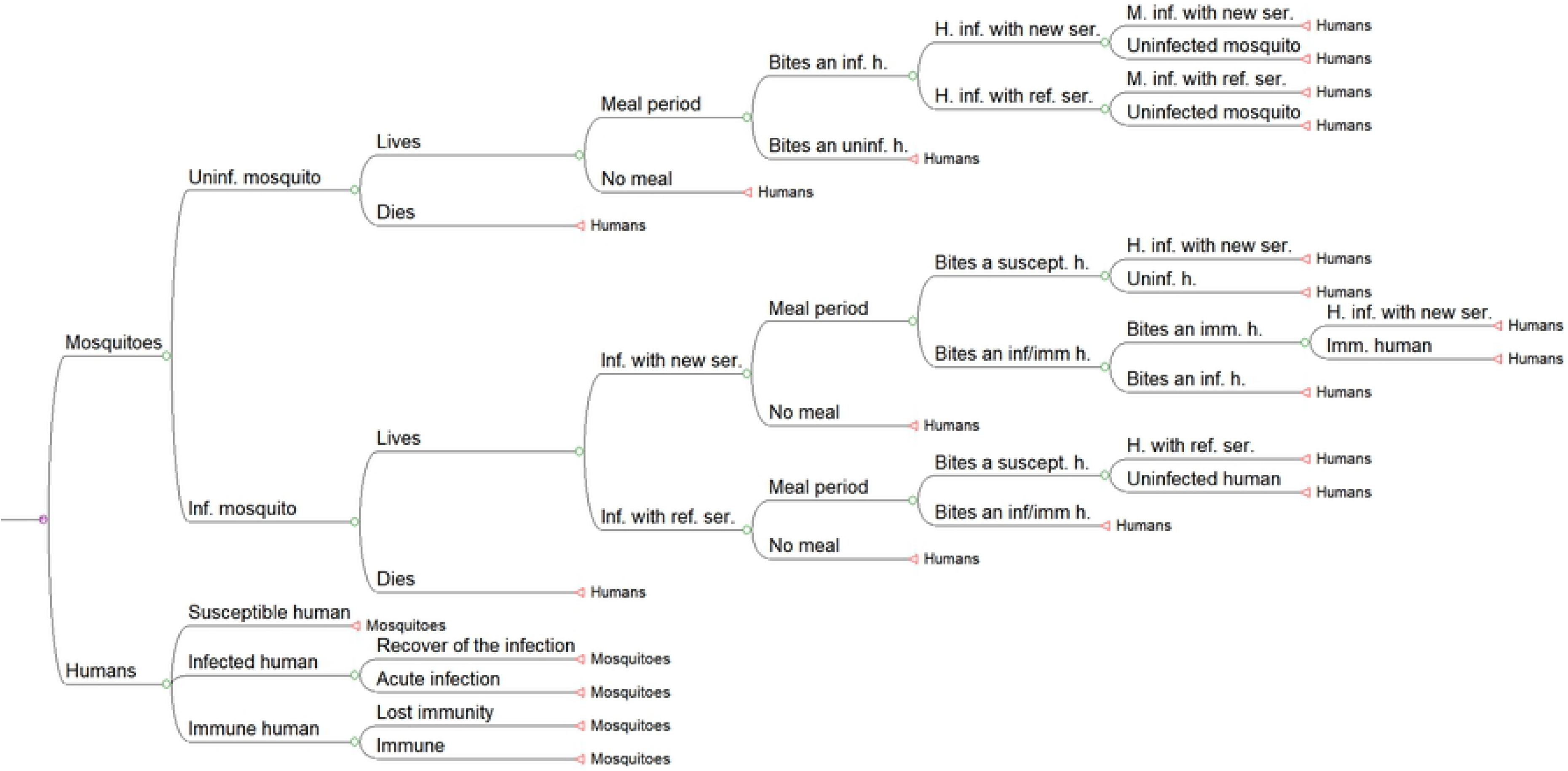
Schematic of the Markov Model. The nodes were the immunological break events and the new serotype establishment events are indicate with arrows. Abbreviations: Inf = Infected, Uninf = Uninfected, H = Human, M = Mosquito, Ref = Reference, Imm = Immune, Ser = Serotype, and Suscept = Susceptible.

The number of infections in humans and mosquitoes are counted, and the probabilities readjusted at each time step. The same occurs with the number of infection recoveries, the TNSEs and the NSCEs. The rain season is modeled as a 4-month period sine function, were the temperature changes from 25 °C to 30 °C, which cross the optimal transmission temperature ranging between 27-28 °C, as we already showed in our previous work (23). The parameters used to calculate the model are show at Table 1. For each mosquito, we implemented an algorithm to introduce mutations in the model. The generation of new serotypes are produced by a stochastic process, were the mutation rate per site is evaluated over the sequence of the E-protein, which is 1.5 Kb long, approximately. To determine if the new sequence corresponds to a new serotype, we established a cutoff of 60% to 80% in the sequence identity. We know from reference (24) that the E-protein has a mutation rate of 1.33×10^-3^ per site/per year, which is then divided by 365. The probability of emergence of new serotypes is obtained by multiplying the mutation rate by the length of the sequence (1.5 Kb) of the E-protein. When the accumulation of mutations reaches 60% - 80% of the protein sequence (the cutoff is selected randomly within this range), the newly generated virus represent a new serotype. The stochastic decision of new serotype emergence is done by a Metropolis–Hastings algorithm (25), if a sequence meet the previous criteria. At this point, we state the “emergence of a new serotype”.

**Table 1.**
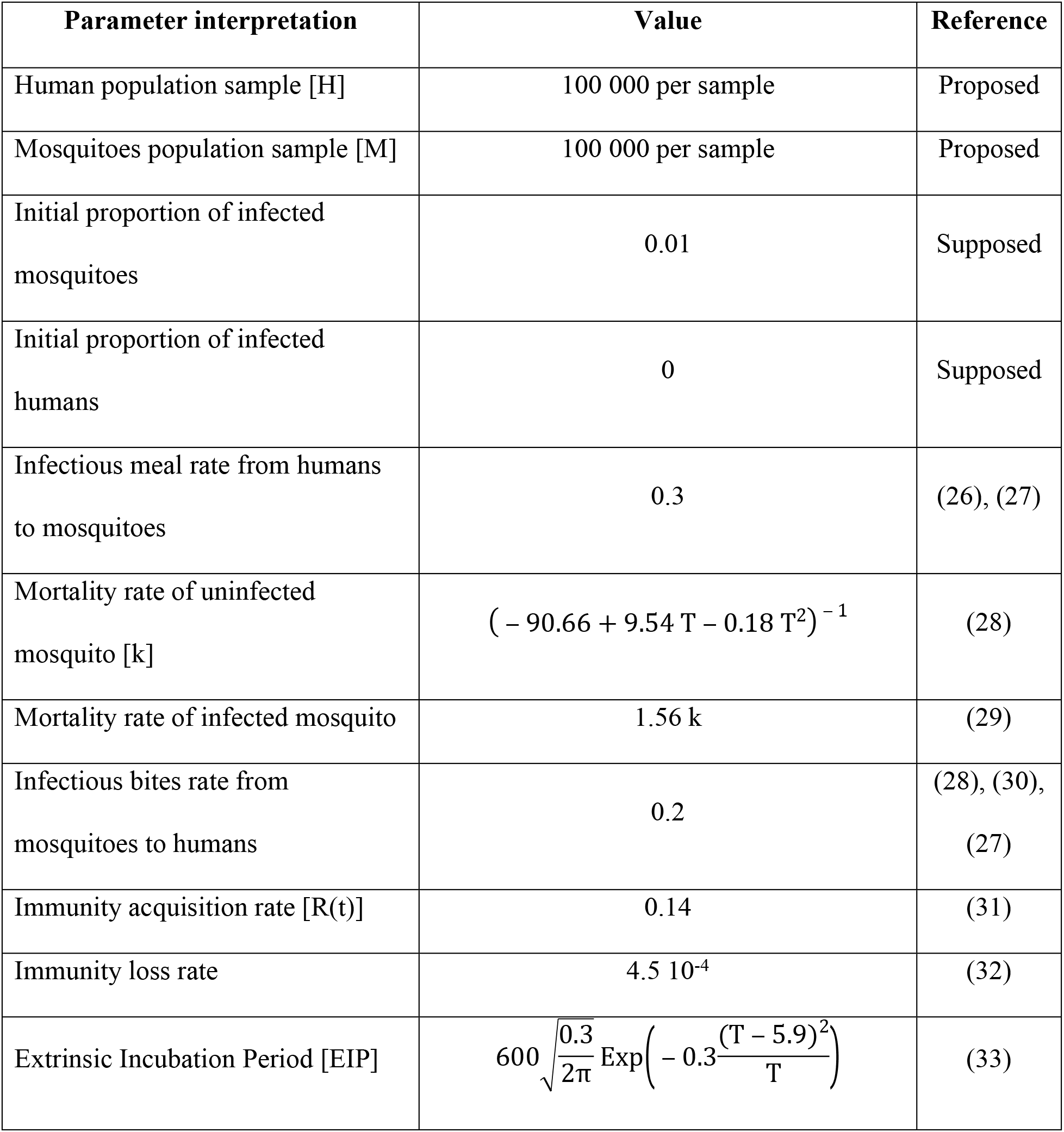
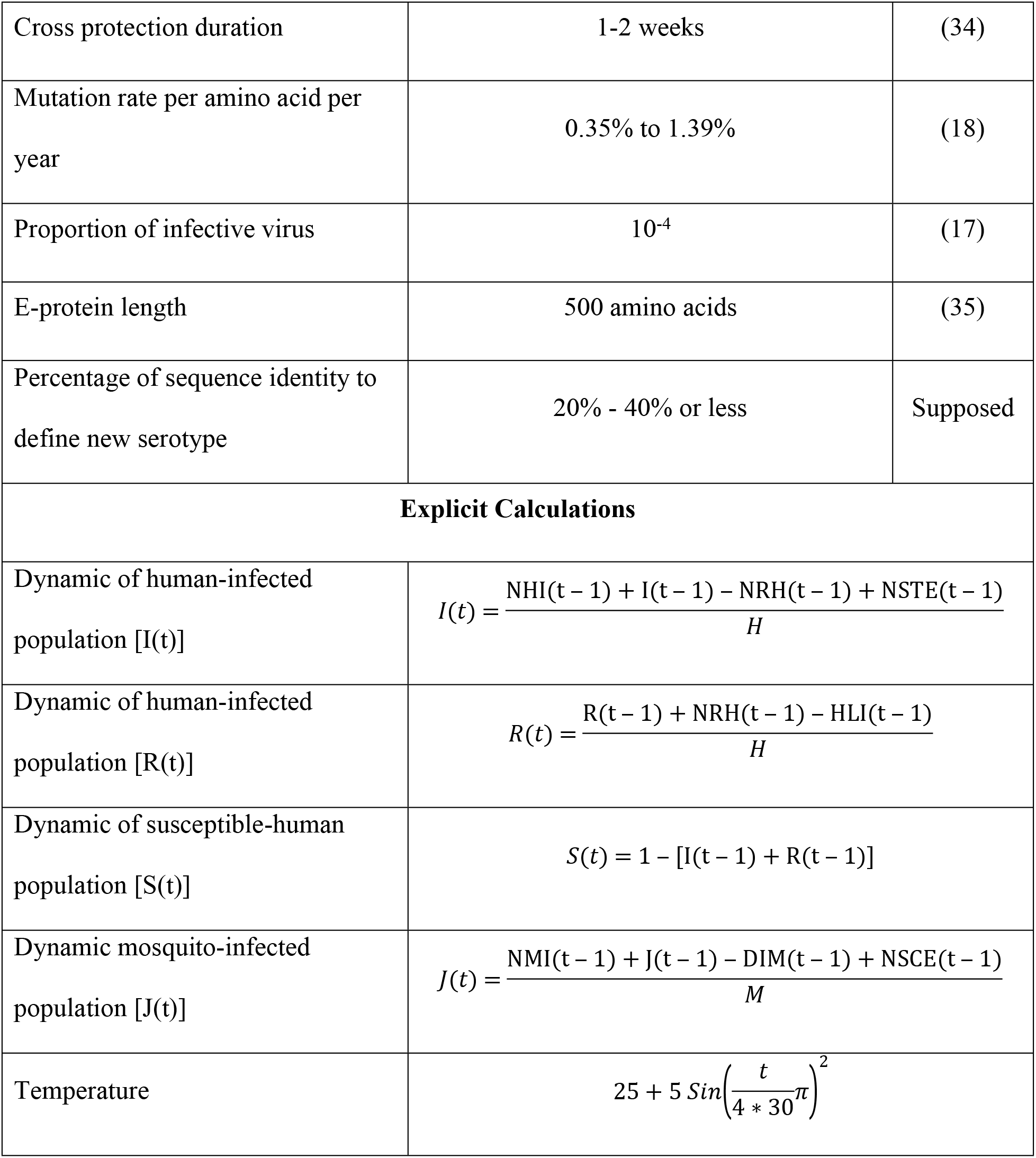
Values and interpretation of the parameters of the model. T = Temperature, t = Time, NSTE = New serotype transmission event, NHI = New human infections, NRH = New recovered human, HLI = Human immunity loss, NMI = New mosquitoes infections, NSCE = New sequence consolidation events, DIM = Death of infected mosquito

Finally, some important points have to be considered for the calculation algorithm: in Table 1, the numerical parameters are used to define probabilistic Beta Distributions, whose mean amounts correspond to the value of the parameter (with 10% standard deviations). Secondly; the parameters represented as functions (for example the EIP function) will be multiplied by 1+ N(0,0.1), were N represents a probabilistic Normal Distributions with zero as mean value and standard deviations of 10%. This factor is a dispersion function that introduces variability to the parameter. The calculations of the model were performed using the TreeAge Pro 2008 program, and the outcomes of the simulations were processed using the standard Microsoft Excel package, only to obtain graphs.

### Scenarios

Multiple scenarios will be solved to investigate the effect of time and vertical transmission on new serotype consolidation in the population. All the scenarios are calculated from 500 samples of the population, with 100 000 persons and 100 000 mosquitoes populations in each sample.

1. Basal scenario. The mutation rate used is the natural rate, with no vertical transmission. The results will be interpreted as the outcomes of the transmission and mutation dynamics over one rain season (one year).
2. The mutation rate is increased 100 times, with no vertical transmission. The results will be interpreted as the outcomes of the transmission and mutation dynamics over 100 rain seasons (100 years).
3. The mutation rate is increased 1 000 times, with no vertical transmission. The results will be interpreted as the outcomes of the transmission and mutation dynamics over 1 000 rain seasons (1 000 years).
4. The mutation rate is fixed as 1 000 times greater than the natural rate, and the proportion of vertical transmission established are 10%, 30% and 50%. The results will be interpreted as the effect of the vertical transmission over the new serotype consolidation.

